# Early life sleep disruption potentiates lasting sex-specific changes in behavior in genetically vulnerable *Shank3* heterozygous autism model mice

**DOI:** 10.1101/2021.09.23.461518

**Authors:** Julia S. Lord, Sean M. Gay, Kathryn M. Harper, Viktoriya D. Nikolova, Kirsten M. Smith, Sheryl S. Moy, Graham H. Diering

## Abstract

**Background:** Patients with autism spectrum disorder (ASD) experience high rates of sleep disruption beginning early in life, however the developmental consequences of this disruption are not understood. We examined sleep behavior and the consequences of sleep disruption in developing mice bearing C-terminal truncation mutation in the high-confidence ASD risk gene *SHANK3* (Shank3ΔC). We hypothesized that sleep disruption may be an early sign of developmental divergence, and that clinically-relevant *Shank3^WT/ΔC^* mice may be at increased risk of lasting deleterious outcomes following early-life sleep disruption.

**Methods:** We recorded sleep behavior in developing *Shank3^ΔC/ΔC^*, *Shank3^WT/ΔC^*, and wild type siblings of both sexes using a non-invasive home cage monitoring system. Separately, litters of *Shank3^WT/ΔC^* and wildtype littermates were exposed to automated mechanical sleep disruption for 7 days prior to weaning (early-life sleep disruption: ELSD) or post-adolescence (PASD) or undisturbed control (CON) conditions. All groups underwent standard behavioral testing as adults.

**Results:** Male and female *Shank3^ΔC/ΔC^* mice slept significantly less than wild type and *Shank3^WT/ΔC^* siblings shortly after weaning, with increasing sleep fragmentation in adolescence, indicating that sleep disruption has a developmental onset in this ASD model. ELSD treatment interacted with genetic vulnerability in *Shank3^WT/ΔC^*mice, resulting in lasting, sex-specific changes in behavior, whereas wildtype siblings were largely resilient to these effects. Male ELSD *Shank3^WT/ΔC^*subjects demonstrated significant changes in sociability, sensory processing, and locomotion, while female ELSD *Shank3^WT/ΔC^* subjects had a significant reduction in risk aversion. CON *Shank3^WT/ΔC^* mice, PASD mice, and all wildtype mice demonstrated typical behavioral responses in most tests.

**Limitations:** This study tested the interaction between developmental sleep disruption and genetic vulnerability using a single ASD mouse model: Shank3ΔC (deletion of exon 21). The broader implications of this work should be supported by additional studies using ASD model mice with distinct genetic vulnerabilities.

**Conclusion:** Our study shows that sleep disruption during sensitive periods of early life interact with underlying genetic vulnerability to drive lasting and sex-specific changes in behavior. As individuals progress through maturation they gain resilience to the lasting effects of sleep disruption. This work highlights developmental sleep disruption as an important vulnerability in ASD susceptibility.

## Background

Sleep is a conserved and essential physiological process that supports neuronal circuit development and lifelong brain health. Up to 80% of individuals with autism spectrum disorder (ASD) suffer from impaired sleep, including difficulties engaging and maintaining sleep (1-3). Moreover, sleep disruption is often seen in advance of ASD diagnosis, and the severity of sleep disruption can be predicative of the severity of other ASD associated phenotypes (4, 5), suggesting that sleep disruption is an early symptom in ASD and a potential driver of the condition. However, it is unclear whether sleep disruption during development, plays a causal role in altered brain maturation, directly contributing to the behavioral symptoms of ASD (6). Recent studies using rodent or fruit fly models showed that early life sleep disruption results in altered cognitive and social behaviors in adults, often with a male bias, strongly suggesting that sleep disruption during development can contribute to lasting ASD relevant phenotypes (7-10).

In humans and other mammals, total sleep amount, and the proportion of rapid eye movement (REM) sleep, is seen at the highest levels directly following birth and during the early postnatal period. The ontogenetic hypothesis of REM suggests that REM sleep produces intrinsic patterns of neural activity that are critical for early postnatal brain development (11). As mammals mature past the early postnatal period, total sleep amount declines and sleep composition transitions from REM sleep prominent to non-REM (NREM) sleep prominent (11). Electroencephalogram (EEG) studies in developing mice show a dramatic remodeling of sleep physiology from postnatal day ∼P10-12 when distinct vigilance states first emerge, to ∼P40 when sleep architecture assumes an adult-like composition and homeostatic response to sleep loss (12, 13). At P10-14 sleep states are highly fragmented, whereas from P14-P21 sleep behavior undergoes considerable consolidation with a rapid increase in bout lengths, decrease in state transitions, and a decrease in the proportion of REM sleep (12, 14). This period also includes important stages of postnatal brain development, including profound synaptogenesis, and maturation of forebrain inhibitory neurons (15-17). However, it remains unclear how the maturation of sleep behavior during early postnatal life contributes to other aspects of brain development and ultimately to adult behavior, and whether disruption of sleep during specific epochs of development leads to specific lasting changes in ASD-relevant behaviors.

*SHANK3* is a high confidence autism risk gene (18) that encodes a prominent excitatory post-synaptic scaffold protein. Heterozygous mutation or chromosomal loss of *SHANK3* is causative of Phelan-McDermid syndrome, a severe neurodevelopmental condition associated with autism and sleep disruption (19-21). ASD model mice with homozygous mutations in *Shank3* exhibit a variety of ASD-relevant phenotypes, while the more clinically relevant genotype, *Shank3* heterozygotes show no or limited phenotypes (22-26), often leading to the use of *Shank3* homozygous mice as a preferred research model. Here we examined developmental sleep behavior and the lasting response to early life sleep disruption (ELSD) in ASD model mice bearing a C-terminal truncation of *Shank3* (*Shank3*ΔC) caused by deletion of exon 21 (24). A recently published study used electroencephalogram (EEG) recordings to show that adult male homozygous *Shank3*^ΔC/ΔC^ mice exhibit clear sleep disruption, reminiscent of patients with Phelan-McDermid syndrome (27). Here, we investigated whether sleep disruption in these mice had a developmental onset. We used a non-invasive method to examine sleep in developing homozygous and heterozygous Shank3ΔC mice of both sexes, together with wild-type littermates and observed that sleep disruption is prevalent *Shank3*^ΔC/ΔC^ males and females as an early phenotype. Developmental onset of sleep disruption was also recently reported in homozygous *Shank3 InsG3680* knock-in ASD model mice, a mutation that affects a similar region of the protein as the Shank3ΔC model used here (10, 28). However, we find the clinically relevant genotype: *Shank3*^WT/ΔC^ heterozygotes, did not show any measured differences in gross sleep parameters compared to WT littermates. Because *Shank3*^WT/ΔC^ heterozygotes do not intrinsically develop overt sleep disruption, these mice are an ideal model to test the potential causal role of developmental sleep disruption in lasting behavior phenotypes, by using an experimentally induced early life sleep disruption method recently established by Jones and colleagues (7, 9). Our results show that a period of early life sleep disruption (postnatal day P14-21), contributes to lasting and sex-specific changes in adult behavior in the genetically vulnerable *Shank3* heterozygous mice. Together, our findings suggest sleep disruption is a phenotype that can emerge early in an ASD mouse model and is an important factor in contributing to lasting changes in brain function and behavior.

## Materials and Methods

### Mice

All animal procedures were approved by the Institutional Animal Care and Use Committee of the University of North Carolina at Chapel Hill and were performed in accordance with the guidelines of the U.S. National Institutes of Health. *Shank3*ΔC mice (deletion of *Shank3* exon 21, described in Kouser et al., 2013(24)) were generously provided by Dr. Paul Worley (Johns Hopkins University) and are maintained on a C57BL/6J background. *Shank3*ΔC mice are commercially available from Jackson Laboratories (strain #018398). Wild-type C57BL/6J breeders were regularly purchased from Jackson laboratories (strain #000664). For sleep recording experiments described in Fig. 1, *Shank3*^WT/ΔC^ heterozygous males and females were crossed to generate cohorts containing WT, *Shank3*^WT/ΔC^ heterozygotes, and *Shank3*^ΔC/ΔC^ homozygotes of both sexes. For the remainder of the experiments (Figs. 2-5), *Shank3*^WT/ΔC^ heterozygous sires were crossed with WT C57BL/6J dams to generate cohorts containing WT and *Shank3*^WT/ΔC^ heterozygotes of both sexes. Following weaning, all experimental mice were group housed with same sex littermates, with the exception of during sleep recordings where mice are individually housed, followed by euthanasia at the end of sleep recording.

**Figure 1.**
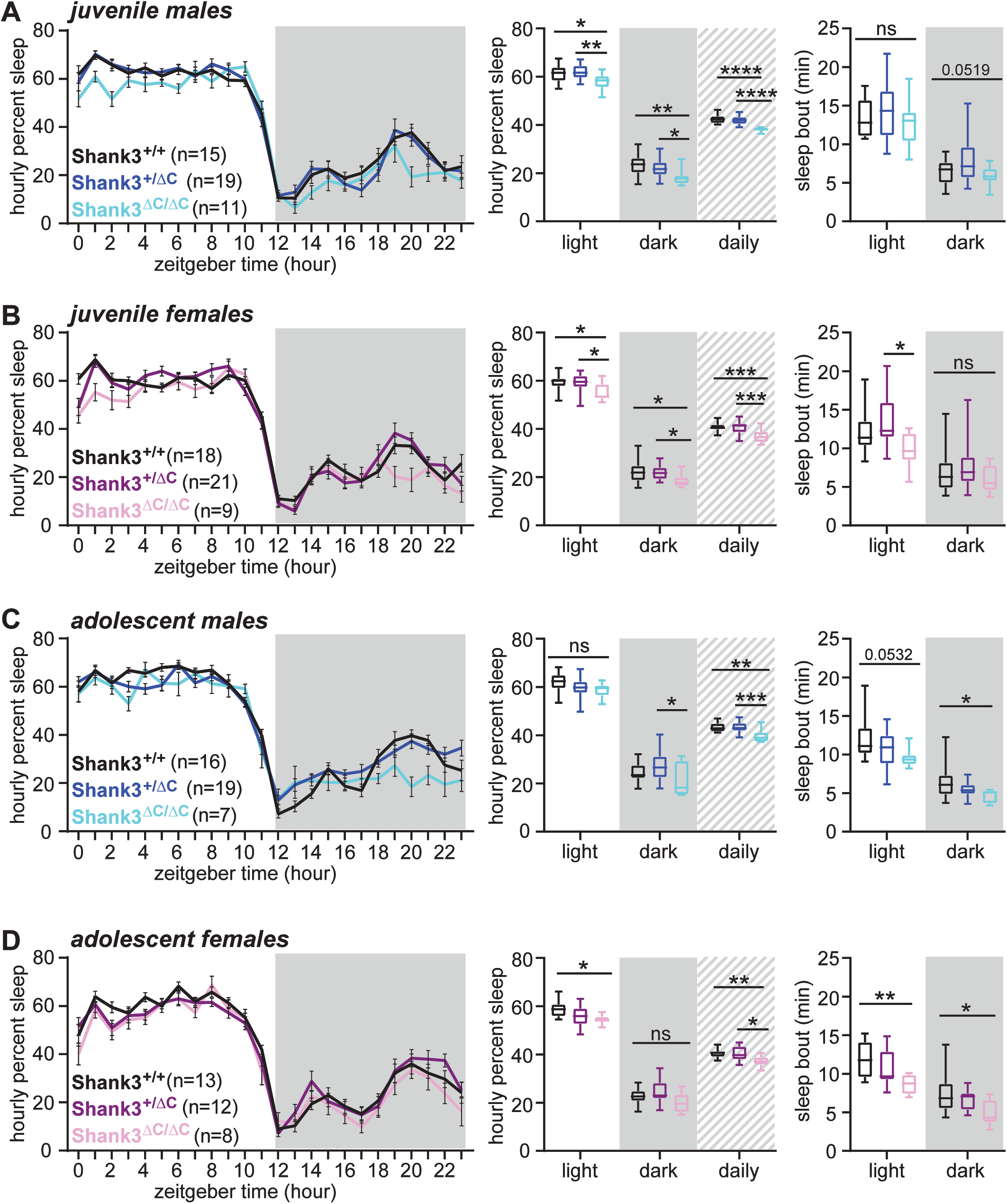
Developmental sleep disruption in *Shank3*ΔC ASD mouse models. (A-D) non-invasive home cage sleep recordings were conducted for male and female *Shank3*^WT/ΔC^ heterozygotes, *Shank3*^ΔC/ΔC^ homozygotes, and wild-type (WT) littermates, at two developmental time points: juvenile (P23-P41), and adolescent (P42-P56). (A) juvenile males, (B) juvenile females, (C) adolescent males, (D) adolescent females. (A-D) Left panel: trace of daily average hourly percent sleep, dark phase indicated by gray box. Center panel: average hourly percent sleep for the light or dark phase (12hrs), or the daily average. Right panel: average sleep bout length observed in the light or dark phase. Genotype differences found with post-hoc Tukey correction are indicated. *P<0.05, **P<0.01, ***P<0.001, ****P<0.0001 (1-way ANOVA with Tukey’s test for multiple comparisons). Error bars indicate ± SEM in sleep traces; box plots show median, range, and 1^st^/3^rd^ quartiles for sleep values. Statistics analysis are summarized in Additional File 1.

**Figure 2.**
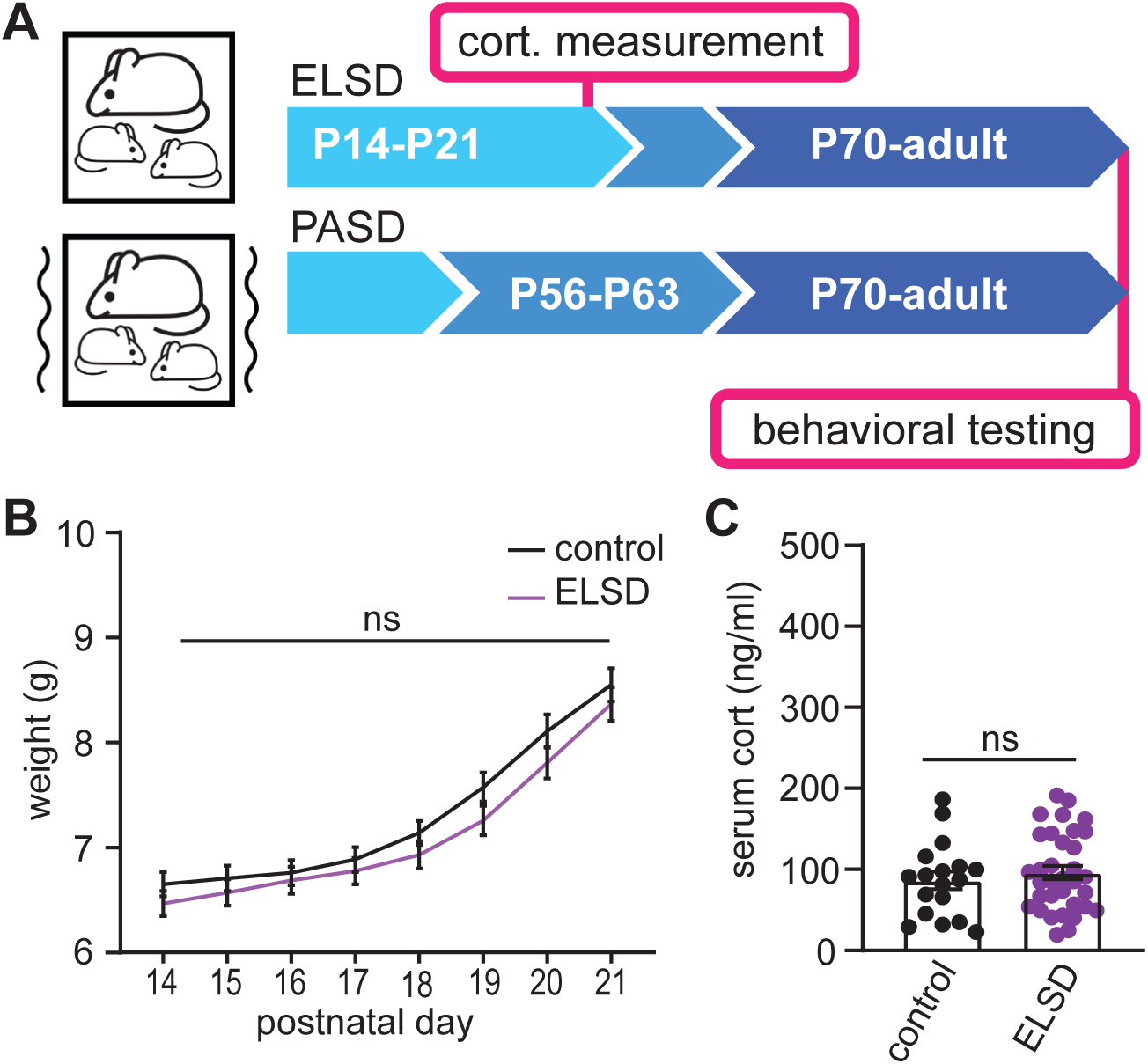
Early life or post adolescent sleep disruption. (A) Experimental design. Cohorts of mice undergo 7 days of sleep disruption either from P14-P21: early life sleep disruption (ELSD), or from P56-P63: post adolescent sleep disruption (PASD). Control cohorts are left undisturbed. Control, ELSD, and PASD cohorts include male and female WT and *Shank3*^WT/ΔC^ heterozygous littermates. Upon reaching adulthood (P70) control, ELSD, and PASD cohorts undergo a panel of behavioral testing lasting 6-7 weeks. (B) Control and ELSD pups were weighed daily from P14-P21. No differences were seen between groups (2-way ANOVA). N=37 control pups; 39 ELSD pups. (C) Separate cohorts of control and ELSD treatment were sacrificed at P21 and serum corticosterone (cort.) was measured using ELISA. No differences were seen between groups (unpaired t-test). N=17 control, 33 ELSD. Error bars indicate ± SEM. See also Additional File 3.

### Sleep behavior recordings

Sleep recordings were conducted in a dedicated sleep behavior room free from any other mouse husbandry activities to allow for uninterrupted behavior assessment, and were maintained on a 12 hours:12 hours light:dark cycle (lights on 7am to 7pm). Sleep/wake behavior was recorded using a non-invasive home-cage monitoring system, PiezoSleep 2.0 (Signal Solutions, Lexington KY). The system uses a piezoelectric mat underneath the cage to detect vibrational movement of the mouse. Customized software (SleepStats, Signal Solutions, Lexington KY) uses an algorithm to process the signal and discern sleeping respiratory patterns from waking respiratory patterns. Sleep was characterized primarily by periodic (2–3 Hz) and regular amplitude signals, which is typical of respiration from a sleeping mouse. In contrast, signals characteristic of wake were both the absence of characteristic sleep signals and higher amplitude, and irregular signals associated with volitional movements, even subtle head movements during quiet wake. The piezoelectric signals in 2-s epochs were classified by a linear discriminant classifier algorithm based on multiple signal variables to assign a binary label of “sleep” or “wake.” Data collected from the cage system were binned over specified time periods: 1hr bins to generate a daily sleep trace, 12hr bins for average light or dark phase percent sleep or sleep bout lengths, 24hr bin for total daily average percent sleep. To eliminate the impact of short and ambiguous arousals on the bout length statistics, a bout length count is initiated when a 30-second interval contains greater than 50% sleep and terminates when a 30-second interval has less than 50% sleep. This algorithm has been validated by using electroencephalography, electromyography, and visual evaluation (29-31). During recording, male and female C57BL/6J wild type, *Shank3*^WT/ΔC^ heterozygous, *Shank3*^ΔC/ΔC^ homozygous mice were individually housed in 15.5 cm^2^ cages with bedding, food and water. Mice were given two full dark cycles to acclimate to the recording cages before the beginning of data collection. No other animals were housed in the room during these experiments. 8-10 days of uninterrupted data were collected for baseline sleep behavior. Separate cohorts were generated for sleep recordings for the juvenile (P25-P41) and adolescent (P42-P56) age groups. 13 juvenile litters were recorded in 6 cohorts, and 10 adolescent litters in 4 cohorts. All animals from the generated litters were recorded and included in analysis minus one juvenile wild type male following equipment failure. Sleep amount and bout lengths were averaged from the multiple days of recordings for each individual prior to statistical comparisons between the experimental groups.

### Early life and post-adolescent sleep disruption (ELSD/PASD)

*Shank3*^WT/ΔC^ heterozygous mice and wild type littermates were subjected to 7 days of 24-hour early life sleep disruption (ELSD, postnatal days 14 – 21) or post adolescence sleep disruption (PASD, postnatal days 56 – 63) using the methods described previously (7). For sleep disruption, the home cage was placed atop orbital shakers and mechanically agitated at 110 rpm for 10 seconds following every 99 seconds still (109 seconds total per cycle), thus producing a mild but frequent mechanical stimulus, shown to interrupt sleep in rodents (32, 33). Control cages were kept atop unplugged orbital shakers and housed in the same room alongside the sleep disruption cohorts. For ELSD experiments each home cage contained 1 entire litter of mouse pups, including littermates of both sexes and genotypes, together with their dam (WT). Eight litters were assigned to each of control and ELSD treatment across five cohorts, with an average of 7.6 pups per litter. Litters were assigned to treatment based on size and sex ratio within their respective cohorts. Prior to start, all pups were confirmed capable of righting themselves immediately from a prone position and initial weights were taken. Individual weights were checked daily during treatment for all pups, and pups were tracked with non-toxic markings on their tails. Body condition of the pups and dams was also assessed daily and judged to be in good appearance. The ELSD treatment was well tolerated, no adverse effects were noted in any pups and no pups were excluded from further analysis. Dams were weighed on the first and last days of treatment. Following treatment, all litters were returned to the vivarium and pups were weaned, genotyped, and socially housed with same-sex littermates. For PASD experiments, young adults were weighed on the first and last days of treatment and visually assessed daily to confirm good body condition. Six litters were used, with an average of 7 mice per litter, and subjects were socially housed in groups of 3 – 5 sex-matched littermates. At the conclusion of the treatment, all animals were returned to the vivarium, prior to further behavioral characterization. Treatment took place in the room described above, with a 12:12 light cycle and no other animals in the room. Subjects were housed in clear standard mouse cages (Tecniplast) with irradiated cobb bedding, cotton nestlet, enrichment hut, and chow *ad libitum*. Hydrogel (ClearH2O) was supplied in place of water bottles during the sleep disruption treatment to prevent splashing and was changed daily.

### Live video recording validation of ELSD

Live-video recording was used to examine wake and sleep behavior in mouse pups aged P14-P21 during control or ELSD treatment conditions. In order to visualize mouse pups, it was necessary to remove the dam, that would otherwise prevent clear observation. In an effort to minimize the interruption to home-cage behavior, we adopted an acclimatization strategy by daily removing the dam from the home cage at the same time of day, ZT4. Four litters of C57BL/6 mice bred in-house were transferred with their dams to a dedicated housing room as described in the ELSD paradigm for acclimatization. Litters were previously culled to 3-4 pups and were randomly assigned to control or ELSD treatment. Each litter was kept in its vivarium home cage and placed atop an unplugged orbital shaker. Initial weights were taken and ears and tails were marked with non-toxic ink. On P12, the dams were removed from the home-cage at ZT4 for 30 minutes to another room and given palatable food, meant to mitigate the stress of handling. The pups remained in the home cage and a heated mat was placed underneath the cage. The cage lid was switched to one customized with a small video camera attached. After 30 minutes, the dam was returned, the heater was turned off, and the cage lid was replaced. On P13-P21 the process was repeated daily at ZT4 with a 60-minute separation period (Extended data Fig. 2-1 D-F). At the conclusion of the separation, weights were taken and ears and tails were re-inked. ELSD treatment was initiated at P14, as described above. Control treatment litters remained on top of an orbital shaker that was left off. Video recording of the pups in their home cage was conducted on P14 and P21. Cage lids were replaced with observation lids and litters were filmed for 60 minutes with Akaso EK70000 Pro cameras at 720p. At the conclusion of the hour, dams were returned, lids were replaced, and weights and inking were repeated. Videos were backed up and converted to MP4 format. Pup ear and tail ink allowed an expert sleep scorer to identify the same individuals at both P14 and P21 time points. A custom scoring template created in The Observer (Noldus) software program described wake and sleep as mutually exclusive behaviors. Only unambiguous wake was scored: myoclonic jerks and swaying due to cage movement or interference from cagemates were not considered wake. Animals were determined to be asleep after 5 seconds of total stillness. Twenty minutes of footage was scored for each individual in each litter, from footage minutes 20 – 40, for an unbiased scoring window. Results were analyzed for mean duration of sleep bouts using GraphPad Prism 9.1.0 (221). At P21, but not at P14, some mouse pups remained awake during the entire 20 min scoring period. We used paired t-tests to compare individual scores from the beginning of the treatment week with scores from the end of the week. Mean group weights at P14 and P21 were analyzed by 2-way ANOVA.

### Serum corticosterone quantification

Control or ELSD treated mice aged P21 were briefly anesthetized with isoflurane, followed by euthanasia by cervical dislocation and decapitation. Trunk blood was collected and mixed with anti-coagulate (5% sodium citrate) to a final concentration of 1% sodium citrate. The blood and anti-coagulate mixture was centrifuged at 2000RCF for 10 minutes. The supernatant (serum) was collected. Corticosterone level in the serum was assessed using the Arbor Assays corticosterone ELISA kit according to the manufacturer’s instructions. Corticosterone concentration is calculated using a colorimetric readout in comparison with a standard curve measured with a BioTek Gen5 plate reader. Analysis was performed with the Arbor Assays DetectX® Corticosterone (OD) software. Results are expressed as ng/ml of corticosterone concentration in mouse serum.

### Behavioral tests

Mice were tested in a standardized battery of assays, conducted in the order listed below, using published methods (34). More stressful procedures (acoustic startle test, buried food test following food deprivation) were carried out near the end of the regimen, to limit confounding effects of repeated testing. Mice were evaluated in only one or two different procedures per week.

#### Elevated plus maze

A 5-min test for anxiety-like behavior was carried out on the plus maze (elevation, 50 cm H; open arms, 30 cm L; closed arms, 30 cm L, walls, 20 cm H). Mice were placed in the center (8 cm x 8 cm) at the beginning of the test. Measures were taken of percent open arm time and open arm entries.

#### Open field

Exploratory activity was evaluated by a 1-hr test in a novel open field chamber (41 cm L, 41 cm W, 30 cm H) crossed by a grid of photobeams (VersaMax system, AccuScan Instruments). Counts were taken of photobeam breaks, with separate measures for locomotor activity (total distance traveled) and vertical rearing movements. Anxiety-like behavior was assessed by measures of time spent in the center region. One wild type ELSD female and one CON *Shank3*^WT/ΔC^ male were excluded from analysis due to a chamber malfunction.

#### Accelerating rotarod

Mice were first given 3 trials on the rotarod (Ugo Basile, Stoelting Co.), with 45 sec between each trial. Two additional trials were conducted 48 hr later, to evaluate consolidation of motor learning. In each trial, rpm (revolutions per minute) progressively increased from 3 to a maximum of 30 rpm. across 5 min (the maximum trial length), and latency to fall from the top of the rotating barrel was recorded.

#### Sociability and social novelty preference in a 3-chamber choice test

Mice were evaluated for social motivation in a rectangular, 3-chambered box (60 cm L, 41. 5 cm W, 20 cm H) fabricated from clear Plexiglass. Dividing walls had doorways allowing access into each chamber. At the start of the test, the mouse was placed in the middle chamber and allowed to explore for a 10 min habituation period, with the doorways into the 2 side chambers open. During the habituation phase, no mice showed a preference for either side of the testing arena. After the habituation period, the test mouse was enclosed in the center compartment of the social test box, and an unfamiliar stranger (a sex-matched, unfamiliar adult C57BL/6J mouse) was placed in one of the side chambers. The stranger mouse was enclosed in a small Plexiglas cage drilled with holes, which allowed nose contact. An identical empty Plexiglas cage was placed in the opposite side of the chamber. Following placement of the stranger and the empty cage, the doors were re-opened, and the subject was allowed to explore the social test box for a 10-min session, concluding the sociability phase of the task. The test mouse was returned to the center chamber and a second stranger mouse was placed in the empty Plexiglass cage. The test mouse was allowed to explore both sides of the chamber for a final 10 minutes, concluding the social novelty preference phase of the task. An automated image tracking system (Noldus Ethovision) provided measures of time spent within 5 cm proximity to each cage, and entries into each side of the social test box.

#### Marble-burying

Mice were tested for exploratory digging in a Plexiglas cage, placed inside a sound-attenuating chamber with ceiling light and fan. The cage floor had 5 cm of corncob bedding, with 20 black glass marbles (14 mm diameter) set up in a 5 X 4 grid on top of the bedding. Measures were taken of the number of marbles buried (i.e., at least two-thirds of the marble covered by bedding) at the end of the 30-min test.

#### Acoustic startle

Mice were evaluated for prepulse inhibition of acoustic startle responses in an automated piezoelectric-based system (SR-Lab, San Diego Instruments). The test had 42 trials (7 of each type): no-stimulus trials, trials with the acoustic startle stimulus (40 msec; 120 dB) alone, and trials in which a prepulse stimulus (20 msec; either 74, 78, 82, 86, or 90 dB) occurred 100 msec before the onset of the startle stimulus. Levels of prepulse inhibition at each prepulse sound level were calculated as 100 - [(response amplitude for prepulse stimulus and startle stimulus together/response amplitude for startle stimulus alone) x 100].

#### Buried food test

Mice were presented with an unfamiliar food (Froot Loops, Kellogg Co.) in the home cage several days before the test. All home cage food was removed 16-24 hr before the test. The assay was conducted in a tub cage (46 cm L, 23.5 cm W, 20 cm H), containing paper chip bedding (3 cm deep). One Froot Loop was buried in the cage bedding, and mice were given 15 min to locate the buried food. Latency to find the food was recorded.

### Statistical analysis

Data were analyzed in GraphPad Prism version 9.1.0 (GraphPad Software, LLC). All results are reported as mean +/- SEM. Sleep characterization was performed with 1-way ANOVA with Tukey’s multiple comparisons test for light/dark cycle sleep amount and bout duration, the results of these tests are summarized in Extended data Table 1-1. Weights were compared using 2-way ANOVA and serum corticosterone was compared using two-tailed unpaired t-test. In the live video validation of the ELSD paradigm, sleep and wake bout lengths during one week of control or ELSD treatment were analyzed using paired t-tests. Behavioral assays were analyzed using multiple unpaired t-tests for genotype with Holm-Šídák method (open field test, elevated plus maze, olfactory, marble burying), multiple paired t-tests for genotype with Holm-Šídák method (sociability), and 2-way ANOVA with Šídák’s multiple comparisons test with genotype and trial as factors (rotarod, pre-pulse inhibition). The results of statistical analysis of mouse behavior testing shown in figure 3 and 4 are summarized in Extended data Table 3-1. Experimental units and n are reported in figures and figure legends. We targeted a minimum sample size of n = 8 per group for ELSD/PASD assays based on recent work by SS Moy and colleagues(34). For sleep recordings, researchers were blind to genotype until data extraction; for sleep disruption, researchers were blind to genotype until the treatment was concluded; for behavioral assays, researchers were blind to genotype and treatment.

**Figure 3.**
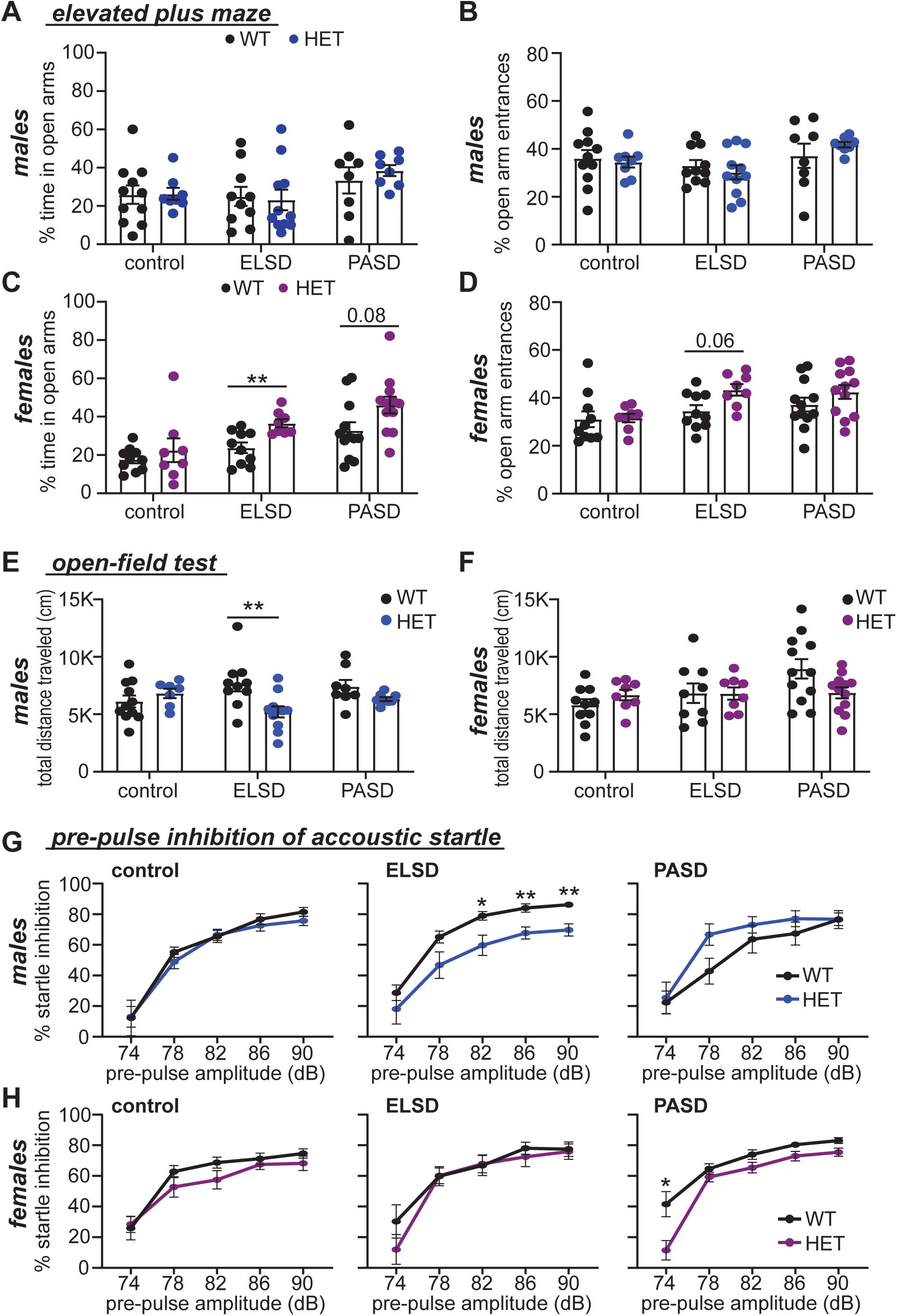
ELSD drives sex-specific changes in certain non-social behaviors in *Shank3*^WT/ΔC^ heterozygotes. (A-D) Percent open arm time and entries for WT and *Shank3*^WT/ΔC^ heterozygotes (HET) from control, ELSD, and PASD treatment groups. (A) Male % time spent in the open arms. (B) Male % open arm entries. No differences were detected in any male groups. (C) Female % time spent in the open arms. ELSD HET females spend significantly more time in the open arms than WT littermates. (D) Female % open arms entries. **P<0.01 (unpaired t-test for genotype with Tukey’s multiple comparisons test). (E-F) Total distance traveled in the open-field task for WT and HET males (E) or females (F). ELSD HET males show hypo-activity in the open field test. (G-H) Percent pre-pulse inhibition (PPI) of acoustic startle responses for WT and HET males (G) or females (H). ELSD HET males show reductions in PPI response. **P<0.01 (Open field, unpaired t-test with Tukey’s multiple comparisons; PPI, 2-way ANOVA with Šídák’s multiple comparisons test). Error bars indicate ± SEM. N=8-12 mice per group/sex/genotype. Summary of statistical analysis is shown in Additional File 2. See also Additional File 4 (anxiety-like measures in open-field test).

**Figure 4.**
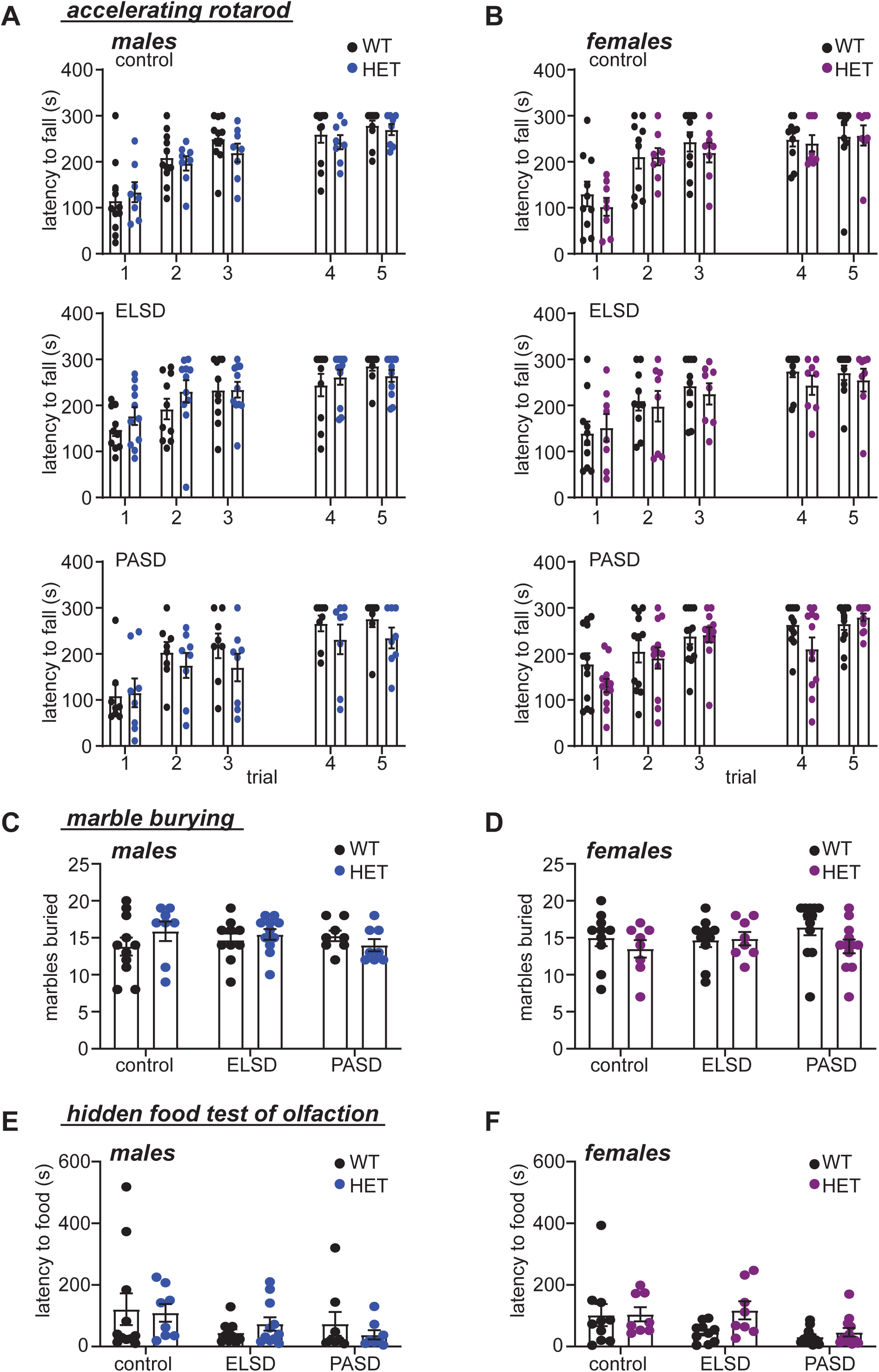
ELSD/PASD treatment did not result in any lasting changes in the accelerating rotarod, marble burying, or hidden food assays. (A-B) Latency to fall in the accelerating rotarod assay of motor coordination and motor learning in control, ELSD, and PASD treated WT and *Shank3*^WT/ΔC^ heterozygous (HET) males (A) and females (B). No differences were observed (2-way ANOVA with Šídák’s multiple comparisons test). (C-D) Marbles buried by control, ELSD, and PASD treated males (C) and females (D). No differences were observed (unpaired t-tests with Holm-Šídák correction). (E-F) Latency to find hidden food in the olfaction test. No differences were observed (unpaired t-tests with Holm-Šídák correction). N=8-12 per treatment/sex/genotype.

## Results

### Sleep phenotypes in male and female *Shank3*ΔC ASD model mice

We first tested whether *Shank3*ΔC ASD model mice exhibited sleep disruption during development. We measured total sleep amount and bout lengths in male and female *Shank3*^WT/ΔC^ heterozygotes, *Shank3*^ΔC/ΔC^ homozygotes, and wild-type (WT) littermates, at two developmental time points: juvenile (P25-P41), and adolescent (P42-P56). Our goals were to determine if developing *Shank3*ΔC exhibit sleep disruption, and to determine when this phenotype emerges. Wake and sleep behavior was monitored for an average of 8-10 uninterrupted days using PiezoSleep, a non-invasive piezoelectric home-cage recording system previously validated using simultaneous EEG/EMG and video recordings (29-31). Average daily total sleep amount and bout lengths were calculated from repeated days of uninterrupted recording for each individual to yield a robust measurement of typical daily sleep behavior. PiezoSleep requires that mice be single housed during recording. Therefore, separate cohorts were generated for the juvenile and adolescent groups in order to mitigate the lasting negative effects of social isolation during development (35). In juveniles and adolescents of both sexes we observed that *Shank3*^ΔC/ΔC^ mice show a significant reduction in total daily sleep compared to WT littermates (1-way ANOVA followed by Tukey’s multiple comparisons test, WT: *Shank3*^ΔC/ΔC^, juvenile males p < 0.0001; juvenile females p = 0.0004; adolescent males p = 0.0021; adolescent females p = 0.009). Reduced sleep amount in juveniles was significant in the light and dark phases (Fig.1A-D). Additionally, in adolescent *Shank3*^ΔC/ΔC^ mice compared to WT, we observed reductions in sleep bout length in the light phase in the female group (female p = 0.0053; male closely approached significance at p=0.0532) and both sexes in the dark phase (male p = 0.0199, female p = 0.0215) (Fig. 1C and D). Post-hoc tests did not reveal any significant differences in sleep amount or bout length between WT littermates and *Shank3*^WT/ΔC^ heterozygotes in either sex or age (Fig. 1A-D, complete reporting of statistical analysis is shown in Additional File 1). Overall, the data show that male and female *Shank3*^ΔC/ΔC^ mice have reduced total sleep detected in juveniles as an early phenotype, and that sleep additionally becomes fragmented as the animals enter adolescence. These findings are in agreement with a recent study describing reduced sleep in developing homozygous Shank3 InsG3680 ASD model mice (10). Additionally, the worsening (fragmentation) of sleep behavior during adolescence in *Shank3*^ΔC/ΔC^ homozygous mice matches observations obtained from some Phelan-McDermid syndrome patients during the transition to adolescence (21, 27, 36). Together with a previous report describing sleep disruption in male adult *Shank3*^ΔC/ΔC^ mice (27), our findings show that sleep disruption in *Shank3*^ΔC/ΔC^ mice has an early onset that persists throughout the lifespan.

### Early life and post-adolescent sleep disruption (ELSD/PASD) in WT and *Shank3*^WT/ΔC^ heterozygotes

In contrast to homozygous *Shank3* ASD mouse models, the clinically relevant genotype, *Shank3* heterozygotes, have repeatedly been shown to have no or limited ASD relevant phenotypes (22-26), in agreement with our sleep measurements (Fig. 1). We speculated that heterozygous mutation of *Shank3* confers a clear genetic vulnerability in mice that may be sensitive to experimental perturbations of sleep during postnatal development. A major question in the ASD field is whether sleep disruption commonly seen in patients, contributes to other ASD phenotypes. Additionally, ASD is diagnosed with an ∼4:1 male bias, although the basis for vulnerability in males or resilience in females is not understood (37, 38). Therefore, we reasoned that *Shank3* heterozygous mice are an ideal model to examine a potential causal role of developmental sleep disruption in lasting sex-specific changes in behavior as a novel gene x sex x environment interaction model for ASD susceptibility.

In order to test whether developmental sleep disruption contributes to altered adult behavior in our *Shank3* ASD model, we utilized the experimental early life sleep disruption (ELSD) paradigm recently established by Jones and colleagues (7, 9). In their recent publications Jones et al., exposed developing prairie voles to ELSD from P14-P21, using an automated method in which the home cage is placed on an orbital shaker, and sleep is regularly disturbed by agitation at 110 RPM for 10 seconds every 109 seconds (10 seconds on, 99 seconds off). In developing voles, this automated method produces a very mild reduction in total sleep amount, but a clear reduction in sleep bout lengths, indicating the method robustly produces sleep fragmentation. Because REM sleep is typically engaged from NREM sleep, one clear effect of this mild but frequent fragmentation method was a significant reduction in total REM sleep (7). ELSD treatment in voles resulted in lasting changes in adult social and cognitive behavior, with a male bias, strongly suggesting that sleep disruption during sensitive periods of brain development is sufficient to drive lasting ASD-relevant phenotypes (7, 9). Similar variations of this automated method have also been shown to cause a reproducible fragmentation of sleep behavior and lasting negative consequences with sustained treatment in adult mice (32, 33, 39, 40).

Here, we tested the lasting effects of this ELSD paradigm on WT and *Shank3*^WT/ΔC^ mice (littermates), reproducing the method described by Jones et al (7, 9)(see methods). To specifically investigate whether early life represents a unique period of vulnerability to sleep disruption, we also tested the effects of sleep disruption treatment delivered later in life from P56-P63, referred to as post-adolescent sleep disruption (PASD) (Fig. 2A). Control groups were placed onto identical orbital shakers that were left off. For ELSD treatments, litters of pre-weaned mouse pups were exposed to sleep disruption from P14-P21, together with their dam (WT) (Fig. 2A). For PASD treatments, separate cages of weaned males and females were exposed to sleep disruption from P56-P63 (Fig. 2A). All groups included WT and *Shank3*^WT/ΔC^ heterozygous littermates of both sexes. Because the dam is also experiencing sleep disruption in the ELSD experiment, pups were examined and weighed daily to ensure that they were receiving sufficient maternal care. Control and ELSD pups maintained healthy appearance and showed comparable daily weight gain (2-way ANOVA: F(1,74) = 1.128, p = 0.2916) (Fig. 2B), and no pups were excluded from further analysis due to any health concerns indicating that the treatment was well tolerated. Interestingly, male *Shank3*^WT/ΔC^ mice in the ELSD group gained slightly more weight than control; however, the weight in this group normalized after weaning and the end of ELSD treatment (Additional File 3). The absence of any weight loss suggests that ELSD treatment did not impair maternal care. To test whether ELSD treatment induces high levels of stress, an additional cohort of mice underwent control or ELSD treatment followed immediately by sacrifice at P21 and measurement of serum corticosterone by ELISA. We did not observe any changes in corticosterone levels between ELSD pups and control (unpaired t-test, t = 0.8765, df = 48, p = 0.3851) (Fig. 2C, and Additional File 3), suggesting that pups acclimate to this treatment without sustained levels of stress. Finally, to confirm that our automated method is able to fragment sleep in developing mice, we used live-video recording to monitor sleep/wake activity of mouse pups undergoing control or ELSD treatment from P14-P21 (see Methods). Our recordings show that mice treated by ELSD show a decrease in sleep bout lengths from P14 to P21 (Paired t-test, control: t(7) = 1.484, p = 0.1814; ELSD: t(6) = 3.923, p = 0.0078) (Additional File 3:D-F). These data suggest that ELSD treatment successfully fragments juvenile sleep but does not overtly impair mouse pup growth and maternal care, or drive a sustained increase in stress, replicating the same conclusions reached from ELSD treatment in prairie voles (7).

Upon reaching P70, control, ELSD and PASD mice underwent an extensive panel of behavioral testing that lasted 6-7 weeks (Fig. 2A). These behavior tests are divided into non-social (Fig. 3-4), and social (Fig. 5); *non-social tests*: elevated plus maze, open-field behavior, pre-pulse inhibition of acoustic startle, accelerating rotarod, olfactory ability, and marble burying; *social*: sociability and social novelty preference in the 3 chamber assay. We asked whether ELSD/PASD treatment caused lasting changes in behavior, whether *Shank3*^WT/ΔC^ heterozygotes were more vulnerable to these treatments than WT siblings, and whether distinct effects were seen between the sexes.

### Sleep disruption during early life contributes to lasting sex-specific changes in specific non-social behaviors in genetically vulnerable *Shank3*^WT/ΔC^ heterozygotes

For the non-social assays, we did not observe any differences in behavior between WT and *Shank3*^WT/ΔC^ littermates in the control treatment group (Fig. 3-4, Additional File 4), consistent with several previous reports (22-26), suggesting that loss of a single copy of *Shank3* alone minimally affects behavior in mice. Strikingly, we observed that ELSD treatment resulted in lasting and sex-specific changes in several non-social behaviors in genetically vulnerable *Shank3*^WT/ΔC^ heterozygotes, while WT littermates were found to be resilient.

ELSD treatment had female specific effect on behavior in the elevated plus maze, an assay of anxiety-like behavior. *Shank3*^WT/ΔC^ females exposed to ELSD treatment showed a significant increase in time spent in the open arms of the plus maze in comparison to WT littermates (unpaired t-test with Holm-Šídák correction: t(16) = 3.51, p = 0.0087), and a similar trend in the number of open arm entries (t(16) = 2.557, p = 0.062). No differences were seen in any male groups (Fig. 3A-D). We describe this female-specific phenotype as “decreased risk aversion”. PASD treatment resulted in a trend of increased time in the open arms of the elevated plus maze (unpaired t-test with Holm-Šídák correction: t(22) = 2.15, p = 0.084) (Fig. 3C), suggesting that *Shank3*^WT/ΔC^ females may have an extended period of vulnerability to the effects of sleep loss in this behavior.

Male-specific effects were seen in open-field behavior and pre-pulse inhibition of acoustic startle response (PPI). In comparison to sex-matched WT littermates, male *Shank3*^WT/ΔC^ heterozygotes exposed to ELSD showed hypo-activity in the open field test as shown by a reduction in total distance traveled (unpaired t-test with Holm-Šídák correction: t(19) = 2.99; p = 0.023) (Fig 3E), whereas no changes were seen in females (Fig. 3F). We did not observe any differences in anxiety-like measures of rearing, or time in the center in the open field test (Additional file 4). Moreover, male *Shank3*^WT/ΔC^ heterozygotes exposed to ELSD showed reduced PPI, a measure of sensory motor gating (2-Way ANOVA: F(1,19) = 5.85, P = 0.026, Main effect of genotype) (Fig. 3G). ELSD treatment showed no effect on PPI behavior in females (Fig. 3H). Interestingly, a small difference in PPI was observed between genotypes in the PASD treated females, albeit at only a single pre-pulse sound (2-way ANOVA: F(1,22) = 0.7215, p = 0.0423 at 74dB, unpaired t-test with Holm-Šídák correction) (Fig. 3H), suggesting a minor change in sensory motor gating.

All groups performed comparably on the accelerating rotarod assay of motor coordination and motor learning, in the marble burying assay for repetitive behavior, and in the hidden-food assay of olfactory ability (Fig. 4). Therefore, decreased risk aversion in ELSD-*Shank3*^WT/ΔC^ females, and hypo-activity in open-field and reduced PPI in ELSD-*Shank3*^WT/ΔC^ males are unlikely to result directly from altered motor behavior, olfaction, or repetitive behavior. In these assays, ELSD treatment had a greater effect on lasting behavior than PASD treatment, suggesting that *Shank3*^WT/ΔC^ mice gain resilience to many of the effects of developmental sleep disruption with maturation (Fig. 3). Together, these results clearly show that ELSD treatment interacts with *Shank3* mutation to potentiate lasting and sex-specific changes in multiple behaviors in genetically vulnerable *Shank3*^WT/ΔC^ heterozygotes.

### Differential effects of ELSD and PASD in social behaviors in genetically vulnerable *Shank3*^WT/ΔC^ heterozygotes

Alterations in social behavior are one of the core diagnostic criteria for ASD, and changes in certain aspects of social behavior are frequently reported in ASD model mice (41). Here we tested the effects of *Shank3*^WT/ΔC^ heterozygous mutation and the effects of our sleep disruption treatments on sociability and social novelty preference using the 3-chamber choice task previously described (26, 42, 43). In the sociability phase of the assay, mice are evaluated for a preference to explore a non-social stimulus (N, a small empty enclosure) or a novel stranger mouse within a small enclosure (social stimulus: S / stranger 1). Social novelty preference is assessed in the subsequent phase by replacing the non-social stimulus with a novel stranger mouse (stranger 2), keeping the now familiar stranger 1 mouse in place. Typically developing mice are expected to spend more time engaging with the social stimulus vs. the non-social stimulus, and to spend more time engaging with novel stranger 2 vs. the already-investigated stranger 1, in the tests for sociability and social novelty preference respectively. Sociability was recently reported to be intact in *Shank3*^WT/ΔC^ heterozygotes of both sexes in comparison to WT littermates, whereas social novelty preference was reported to be impaired in both male and female *Shank3*^WT/ΔC^ heterozygotes (26). Moreover, a recent publication has shown that a period of adolescent sleep disruption can impair social novelty preference in WT C57BL/6J mice, while having no effect on sociability (10). Therefore, these recent findings, together with previous work, have shown that sociability and social novelty preference are clearly distinct phenotypes, with different underlying neural circuits and separate genetic and environmental vulnerabilities (41).

Under control conditions, we observed the expected social approach behavior in WT mice of both sexes (42). As previously reported, under control conditions *Shank3*^WT/ΔC^ heterozygotes of both sexes showed intact sociability, comparable to WT littermates (Fig. 5A-C) (26, 44). In response to ELSD or PASD treatments, all groups showed intact sociability, with the clear exception of male ELSD-*Shank3*^WT/ΔC^ heterozygotes that exhibited a lack of preference for the social stimulus (S) in comparison to the non-social stimulus (N) (ELSD HET p = 0.125; ELSD WT p = 0.0006; CON HET p = 0.0384; CON WT p = 0.02, paired t-test with Holm-Šídák correction for all social behavior results) (Fig. 5A-C). Importantly, sociability was intact in male PASD-*Shank3*^WT/ΔC^ heterozygotes. These findings show that sociability is uniquely affected in genetically vulnerable males exposed to developmental sleep disruption, and that these individuals gain resilience to these negative effects with maturation. (Complete reporting of statistical analysis for social assays is found in Additional File 2.)

The subsequent test for social novelty preference was found to be influenced by sex, genotype, and PASD treatment, but not ELSD treatment. WT males showed intact social novelty preference under all conditions. Interestingly, where male ELSD-*Shank3*^WT/ΔC^ heterozygotes showed impaired sociability (Fig. 5B), these same individuals displayed fully intact social novelty preference (ELSD HET p = 0.0003; ELSD WT p = 0.006) (Fig. 5D). In male PASD-*Shank3*^WT/ΔC^ heterozygotes sociability was intact, but social novelty preference was impaired (Fig. 5D), albeit, these individuals still showed a trend towards preference for the novel stranger 2 (PASD HET p = 0.085; PASD WT p = 0.011), suggesting that social novelty behavior in genetically vulnerable males is sensitive to sleep disruption at later ages of development, consistent with recent reporting (10). For the female groups, in our hands, we observed that only female *Shank3*^WT/ΔC^ heterozygotes showed impaired social novelty preference under control conditions (Fig. 5D-E). This is in contrast to a recent publication reporting impaired social novelty preference in *Shank3*^WT/ΔC^ heterozygotes of both sexes (26). WT female littermates showed expected social novelty preference under control and ELSD conditions, but this behavior was found to be impaired in response to PASD treatment (PASD WT p = 0.298) (Fig. 5E), consistent with a vulnerability in this behavior to adolescent sleep loss (10). Thus, in our hands, social novelty preference was found to be affected by sex, genotype, and PASD treatment. *Shank3*^WT/ΔC^ females were the most vulnerable, showing baseline impairments. WT females and *Shank3*^WT/ΔC^ males showed some selective vulnerability to PASD treatment. WT males were resilient to the treatments tested here. Overall, our findings show that developmental sleep disruption interacts with underlying genetic vulnerability and sex in *Shank3*^WT/ΔC^ heterozygotes to drive lasting and sex-specific changes in specific aspects of non-social and social behaviors.

**Figure 5.**
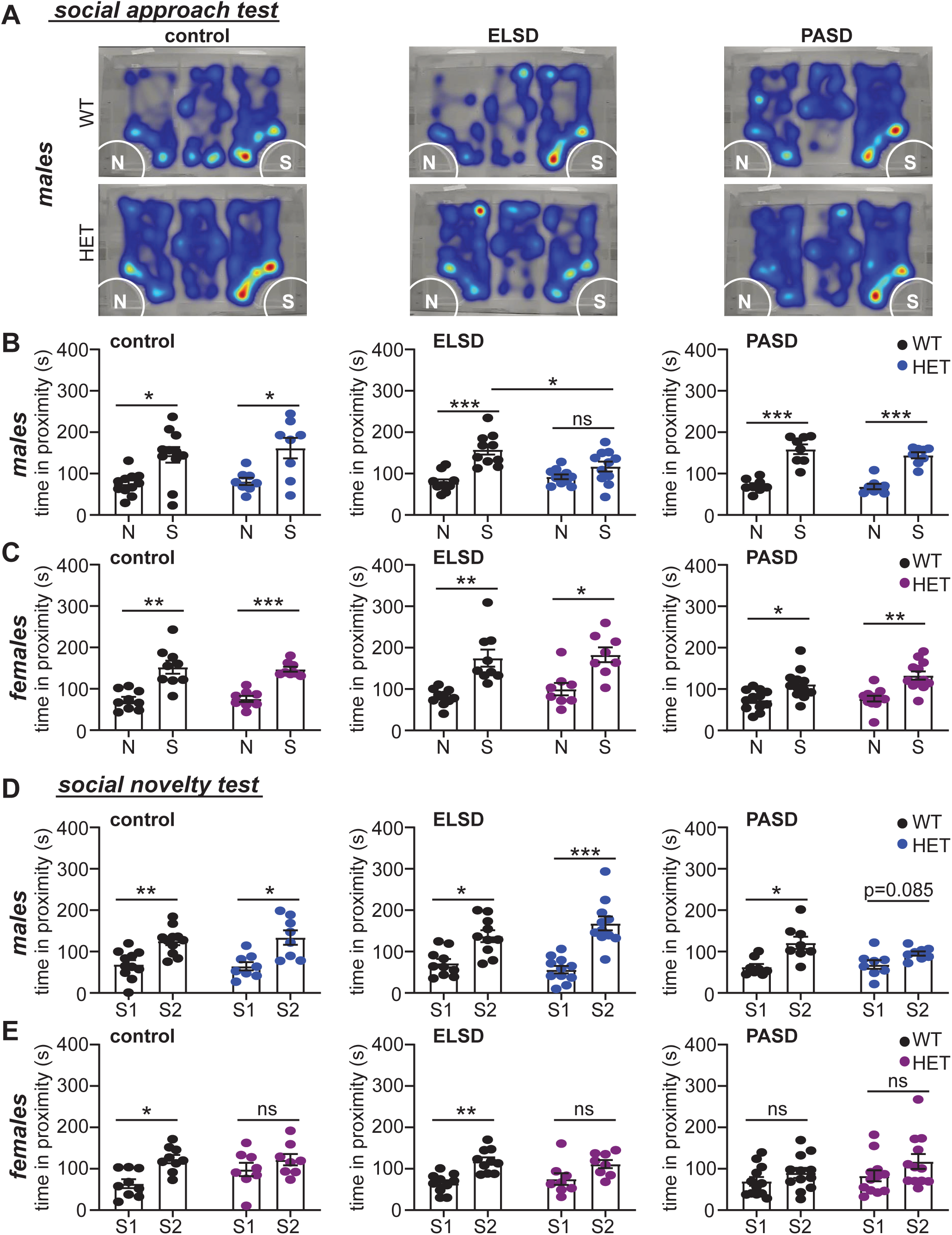
Social approach and social novelty preference behaviors are differentially sensitive to sex, Shank3 genotype, and sleep disruption treatments. (A) Examples of individual WT and *Shank3*^WT/ΔC^ heterozygotes (HET) from control, ELSD, and PASD treatment groups performing the 3-chamber social approach task. Heat map indicates time in location. Mice can freely move between 3-chambers and choose to spend time engaging with the non-social stimulus (N) or with a social stimulus (S). (B-C) Social approach behavior: time spent in proximity with the non-social (N) and social stimulus (S) for WT and HET males (B) or females (C). ELSD HET males show no preference for the social stimulus. (D-E) Social novelty preference: time spent in proximity with familiar stranger 1 (S1) and novel stranger 2 (S2) for WT and HET males (D) or females (E). PASD HET males and WT females show no preference for S2. HET females show now preference for S2 under any condition tested. *P<0.05, **P<0.01, **P<0.001 (paired t-test with Holm-Šídák correction). N=8-12 per treatment/sex/genotype. Summary of statistical analysis is shown in Additional File 2.

## Discussion

Sleep disruption is a common comorbidity affecting a majority of individuals with ASD, including Phelan-McDermid syndrome (1, 2, 20, 27). Problems with sleep are often apparent in human patients prior to an ASD diagnosis, suggesting that sleep disruption may be an early phenotype in the ASD population (5). Sleep disruption has been reported in a number of mouse models of human neurodevelopmental disorders and ASD, including models of fragile X, Rett, Angelman, and Phelan McDermid syndromes (27, 45-47). Whether sleep disruption emerges as a developmental phenotype in mouse models of neurodevelopmental disorders is largely unexplored. In a contemporary study, Medina et al., (BioRxiv, https://doi.org/10.1101/2021.03.10.434728) used EEG recording to obtain longitudinal measurements of sleep behavior in developing homozygous *Shank3*^ΔC/ΔC^ males and WT littermates. This study showed that beginning as early as could be measured at P23, homozygous *Shank3*^ΔC/ΔC^ males sleep less than WT littermates and show evidence of sleep fragmentation, phenotypes that the *Shank3*^ΔC/ΔC^ males maintain throughout their lifespan (27). Here, using an alternative, non-invasive method of sleep analysis we qualitatively replicate these results and reach identical conclusions: sleep disruption emerges as a developmental phenotype in *Shank3*^ΔC/ΔC^ homozygotes. We extend this conclusion by demonstrating a similar developmental sleep disruption in *Shank3*^ΔC/ΔC^ females. Our results are also consistent with the developmental sleep disruption observed in a similar homozygous *Shank3* mouse model InsG3680 recorded from P35-P42 (10). Together, these studies and our data show that sleep disruption emerges as an early phenotype in Shank3 homozygous ASD mouse models of both sexes. Based on the detection of sleep disruption in *Shank3* homozygotes at the start of recording in our current study and in contemporary studies, it is likely that sleep disruption phenotypes emerge at early stages even prior to weaning. This raises a critical question: does developmental sleep disruption play a causal role in altered brain function and behavior in ASD?

ASD-relevant behavioral phenotypes have been repeatedly reported in a number of *Shank3* homozygous mouse models, whereas these phenotypes are largely absent or considerably milder in the clinically relevant genotype: *Shank3* heterozygotes (22-27). Accordingly, in our own hands, we did not detect any significant differences in total sleep amount or bout lengths between *Shank3*^WT/ΔC^ heterozygotes and sex-matched WT littermates at any age tested (Fig. 1). Does the developmental sleep disruption intrinsic to the more severe *Shank3* homozygous ASD models play a causal role in their ASD-relevant adult behavioral phenotypes? Excitingly, a recent study from Bian and colleagues (10) showed that improving sleep during a sensitive period of adolescent development in *Shank3* homozygous InsG3680 mice through pharmacological or optogenetic means, indeed rescued adult deficits in social behavior. Here we show that experimentally induced early life sleep disruption results in lasting and sex-specific changes in behavior in clinically relevant *Shank3* heterozygous mice. The current findings build on previous publications that have shown that early life sleep disruption drives altered adult behavior in drosophila and rodent models (7-9). These studies, together with our current findings, strongly suggest that sleep disruption during sensitive periods of brain development can indeed cause lasting changes in brain function and behavior, particularly with underlying genetic vulnerability, supporting that developmental sleep disruption in ASD patients contributes to other behavior phenotypes in this condition.

### Early life sleep, and brain development

In humans and other mammals, sleep behavior and composition are highly dynamic during postnatal development, where REM sleep is seen in its highest proportions in the early postnatal period, and slow wave sleep (SWS) peaks during adolescence (10, 12, 13, 48). It is plausible that transitions in sleep physiology during development support different epochs of brain maturation. If this is indeed the case, then experimental (or intrinsic) disruption of sleep at specific postnatal ages may result in distinct effects on adult behavior; similarly, the severity of sleep disruption may drive differential consequences. A prominent theory in the field suggests that high levels of REM sleep during early postnatal life produces intrinsic neuronal activity that is essential for healthy brain development (11). In particular, REM sleep during development is believed to be critical for the formation, plasticity, and pruning of neuronal synapses (49-51). Here, we reproduced as closely as possible, the P14-P21 ELSD paradigm used by Jones and colleagues, who showed that this treatment causes sleep fragmentation and consequent selective loss of total REM sleep, resulting in a long-lasting change in partner preference in male, but not female voles, an important social behavior in this species (7). A subsequent study using this paradigm showed that ELSD treatment impairs cognitive flexibility in prairie voles of both sexes, clearly showing that both sexes are vulnerable to developmental sleep loss (9). Replicating the ELSD experimental paradigm using a mouse model, our results qualitatively confirm the conclusions from Jones et al., that early life sleep disruption contributes to lasting changes in behavior in both sexes, including sociability in males (7) (Fig. 5). By using the clinically relevant *Shank3*^WT/ΔC^ heterozygous model, we extend the findings of Jones et al., to demonstrate that developmental sleep disruption interacts with underlying genetic vulnerability. Due to the technical challenges of performing longitudinal EEG recordings from pre-weaned mouse pups (12), we are not able to confirm that this treatment has a selective effect on REM sleep in mouse pups. However, we did confirm using live-video scoring that the paradigm produces sleep fragmentation as reported (7). We additionally replicate the previous findings that this ELSD treatment does not impair pup growth or drive any sustained increases in stress, ruling out changes in stress or nutrition as drivers of lasting behavioral changes. In our hands, wild-type C57BL/6J mice were resilient to the effects of this ELSD paradigm, unlike the wild-derived voles used by Jones et al. This difference may be due species-specific vulnerability, to the greater genetic diversity in the wild-derived voles compared to inbred C57BL/6J lab mice, or the difference in how sociability was measured. Future research using this ELSD paradigm in combination with different lab mouse strains, and in additional genetically vulnerable mouse ASD models will be needed to identify further genetic interactions with developmental sleep loss.

At present, we are unable to determine whether a parsimonious mechanism(s) underlies the various and sex-specific behavioral phenotypes described in this study. The P14-21 ELSD window targeted here represents an important time of maturation of sleep behavior, and other aspects of brain development including a profound period of synaptogenesis and maturation of inhibitory circuits in the cortex (12, 15-17). Changes in synapse density, morphology, and function have been documented repeatedly in animal ASD models and post-mortem patient samples, leading many to describe ASD as a synaptopathy, a disease of the synapse (52-54). This is further supported by the discovery of high confidence ASD risk genes, many of which encode for prominent synaptic proteins, including *SHANK3* (18, 53, 55). Moreover, sleep disruption during development has been shown to affect cortical synapse formation, growth, plasticity, and pruning (49-51). We speculate that sleep disruption during this period interacts with underlying genetic vulnerability, negatively impacting synapse maturation in sensitive neural circuits. In support of this idea, Jones at al., have shown that ELSD treatment in prairie voles drives increased dendritic spine density of layer II/III pyramidal neurons in the pre-frontal cortex (9), indicating a lasting change in glutamatergic synapses. Whether similar changes are seen in *Shank3*^WT/ΔC^ heterozygous mice exposed to ELSD will need to be determined. We speculate that Shank3 may be engaged in protective or homeostatic adaptations, as has been suggested by prior work (44, 56), that mitigate the effects of ELSD treatment in WT mice, thus reduced Shank3 function results in vulnerability to sleep loss in *Shank3*^WT/ΔC^ heterozygotes.

### Adolescent sleep and social novelty preference

In this study we also treated our mice with a later period of sleep disruption from P56-P63, referred to as post-adolescent sleep disruption (PASD). In each case where we observed an effect of the ELSD treatment in Shank3 heterozygotes, we found that PASD treatment did not have the same effect (Fig. 3, and male social approach in Fig. 5A and B), consistent with the idea that early development is uniquely vulnerable to sleep disruption. Interestingly, we found that PASD treatment appeared to impair social novelty preference in *Shank3*^WT/ΔC^ heterozygous males, and WT females, whereas *Shank3*^WT/ΔC^ heterozygous females showed impaired social novelty preference at baseline. We believe these findings are in general agreement with the recent study of Bian and colleagues, reporting that a period of adolescent sleep disruption results in lasting changes in social novelty preference (10). These authors used a paradigm of “adolescent” sleep disruption targeting ages P35-P42 (4hrs near total sleep loss per day, 5 consecutive days within the P35-P42 window), resulting in impaired social novelty preference, but not sociability, in adult wild-type C57BL/6J mice. Sleep disruption from P42-P49 also affected social novelty preference, but to a lesser degree, whereas the same sleep disruption treatment in adults from P84-91 had no effect, leading these authors to similarly conclude that individuals gain resilience to sleep loss with maturation (10). This recent study did not test the P56-P63 “PASD” window used here. Our results suggest that WT C57BL/6J males are already resilient to sleep loss by this age, whereas WT females and *Shank3*^WT/ΔC^ heterozygous males have an extended period of vulnerability to sleep loss, where our PASD treatment resulted in impaired social novelty preference. A body of work shows that sociability and social novelty preference are mediated by distinct neuronal circuits, with differential genetic and environmental vulnerabilities (10, 41). Therefore, it is possible that sleep disruption during the earlier period (P14-P21) negatively impacts the development of circuits underlying sociability in vulnerable males, whereas the later intervention (P35-42;P42-49; P56-63) has a more selective effect on social novelty preference. Bian et al., show that adolescent sleep disruption alters the activation and release patterns of dopaminergic neurons of the ventral tegmental area (VTA). Future studies using sleep disruption interventions at different postnatal ages, and using different mouse ASD models or genetic backgrounds will be important to map the epochs of vulnerability to developmental sleep loss, the specific behavioral consequences, as well as the underlying genetic and circuit vulnerabilities.

### Sex-specific vulnerabilities in ASD

ASD is diagnosed with a clear male bias of ∼4:1, based on the core features of altered social interaction, communication deficits, and abnormal restricted/repetitive behavior. The basis of vulnerability in males, or resilience in females, that underlies this clear sex bias is not known (38). However, it has been suggested that the genetic and environmental vulnerabilities underlying ASD susceptibility may in fact affect males and females in equal numbers, but that the phenotypic presentations diverge, particularly for social behaviors. Consequently, many affected females escape ASD diagnosis, as the diagnostic criteria are based on typical male presentation (37, 57, 58). Unfortunately, this may lead to a lack of treatment in affected females. Our results show that *Shank3* mutation and developmental sleep disruption interact to drive sex-specific changes in behavior, supporting the idea that both sexes share vulnerability, but diverge in the phenotypic outcomes. In particular, our ELSD treatment resulted in altered sociability in *Shank3* heterozygous males, and decreased risk aversion in *Shank3* heterozygous females, clearly showing that both sexes have vulnerability to developmental sleep disruption. Overall, this work points towards early life sleep as a risk factor in ASD susceptibility in both sexes. Future work in this area will be needed to uncover the mechanistic basis by which early life sleep disruption impacts lasting and sex-specific changes in behavior.

## Limitations

Sleep disruption is a common comorbidity in the ASD population but the role of sleep disruption in the condition is unclear (6). Sleep disruption is also seen in patients with Phelan McDermid Syndrome (PMS), known to be caused by heterozygous mutation or chromosomal loss of the *SHANK3* gene (18, 20, 27), and in non-human primates with heterozygous *SHANK3* mutation (59). In contrast to humans and primates, heterozygous mutation of *Shank3* in mouse models results in no or limited behavioral phenotypes, and no gross sleep disruption, suggesting that mice are more resistant to *Shank3* loss of function than humans. Here we show that experimentally induced sleep disruption during development interacts with underlying genetic vulnerability in *Shank3*^WT/ΔC^ heterozygous ASD model mice to drive lasting changes in behavior, demonstrating a causal relationship between developmental sleep disruption altered behavior in this ASD mouse model. Considerably more research will be needed to establish whether a similar causal relationship between sleep disruption and lasting behavioral phenotypes exists in human PMS/ASD patients, where *SHANK3* mutation has a more detrimental effect on brain function. Moreover, the mouse model used here has a deletion of exon 21 of the mouse *Shank3* gene, resulting in expression of a C-terminal truncation (ΔC) (24). Future work should be done to examine whether sleep disruption during development also contributes to lasting changes in behavior in distinct ASD animal models (60), and in alternative *Shank3* mouse models that target different exons within the mouse *Shank3* gene, including the recently developed “complete *Shank3* deletion” targeting exons 4-22, that better models the chromosomal loss of *SHANK3* seen in the majority of PMS patients (25). In our first experiment we used a non-invasive sleep monitoring system to show that *Shank3*^ΔC/ΔC^ homozygotes show a reduction in total sleep time and sleep fragmentation as adolescents, whereas *Shank3*^WT/ΔC^ heterozygotes showed no difference from WT littermates. With our sleep recording system, we are not able to make any conclusions regarding sleep architecture (NREM and REM sleep) or spectral power. It is possible that a more detailed analysis of the sleep phenotypes using EEG recording will reveal subtler sleep phenotypes in *Shank3*^WT/ΔC^ heterozygotes. Related to this, we are not able to determine whether the sleep fragmentation paradigm used here is able to cause a selective loss of REM sleep, as has been previously reported for prairie voles (7).

## Conclusions

Here we show that early life sleep disruption interacts with genetic vulnerability in *Shank3*^WT/ΔC^ heterozygous ASD model mice to drive lasting and sex-specific changes in behavior. This work adds to a growing body of work using drosophila, prairie voles, and mice, showing that sleep disruption during vulnerable periods of brain development drive long-lasting changes in brain function and behavior (7-10). The broader implications of this work are that developmental sleep disruption commonly seen in ASD patients may contribute to other phenotypes in the condition. This work places a greater emphasis in understanding and treating sleep disruption in ASD and other neurodevelopmental conditions. Clearly, more research in this area is warranted.

## Declarations

### Conflict of interests

The authors declare no competing interests

### Funding

This work was supported by a Bridge to Independence Award to GHD from the Simons Foundation Autism Research Initiative (SFARI), and by the Mouse Behavioral Phenotyping Core of the UNC Intellectual and Developmental Disabilities Research Center (NICHD; P50 HD103573; PI: Joseph Piven).

### Author contributions

JSL and SMG generated and genotyped all the mice for this study and performed sleep recording analysis, and participated in intellectual design of the project, analysis of the results and preparation of the manuscript and figure. KMH and VDN performed the behavior assays and participated in data analysis. KMS performed the ELISA measurements of serum corticosterone and participated in data analysis. SSM and GHD designed and supervised the project, prepared the figures, and wrote the manuscript.

## Acknowledgements

We wish to thank Dr. Paul Worley (Johns Hopkins University) for generously providing the *Shank3*ΔC ASD mouse model. We wish to thank Dr. Carolyn Jones-Tinsley and Dr. Miranda Lim (Portland VAMC and Oregon Health Sciences University) for consultation and technical advice on establishing the early life sleep disruption paradigm based on their recent publications (7, 9).

## Additional Files

Additional File 1. Summary of ANOVA statistics for sleep recordings in Figure 1.

Additional File 2. Summary of statistics for behavior tests in Figures 3 and 5.

**Additional File 3.**
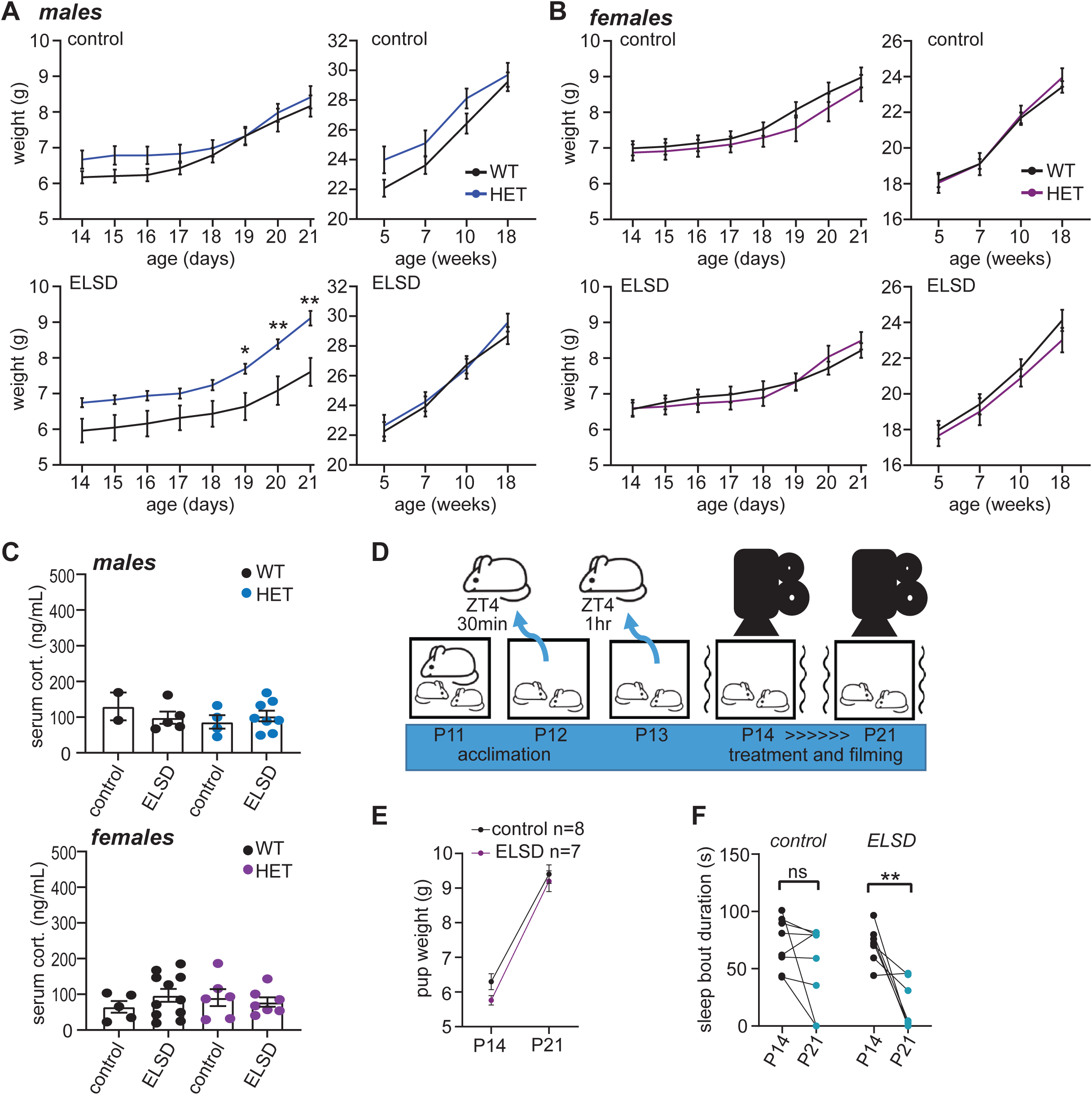
Weight gain, corticosterone levels, and in-treatment sleep bouts in control and ELSD cohorts. (A-B) Control and ELSD cohorts were weighed daily from P14-P21 to ensure that pups were receiving sufficient maternal care during ELSD treatment, and at multiple time points during the remainder of the experiment. (A) weight of male control and ELSD cohorts. *Shank3*^WT/ΔC^ heterozygotes (HET) exposed to ELSD gain weight faster than WT littermates (2-way ANOVA: main effect of genotype: F(1,19) = 6.766, p = 0.175); however, this difference was normalized by 5 weeks of age. N=8-11 per treatment/genotype. *P<0.05, **P<0.01 (2-way ANOVA). (B) Weight of female control and ELSD cohorts. No differences in weight between treatments or genotypes was observed (2-way ANOVA). N=8-12 per treatment/genotype. (C) Serum corticosterone (cort.) was measured using ELISA from separate cohorts of control and ELSD treated pups at P21. No differences in cort. levels between treatments or genotypes within each sex were observed (1-way ANOVA). (D-F) two ELSD and two CON litters were filmed to confirm sleep disruption; (D) experimental design. Pups were acclimated to dam removal at ZT4 prior to live-video recording on P14 and P21. (E) ELSD did not affect weight gain. (F) Seven days of ELSD treatment significantly reduced sleep bout duration by postnatal day 21 **P<0.01 (paired t-test, control: t(7) = 1.484; ELSD: t(6) = 3.923). N=7-8 per treatment.

**Additional File 4.**
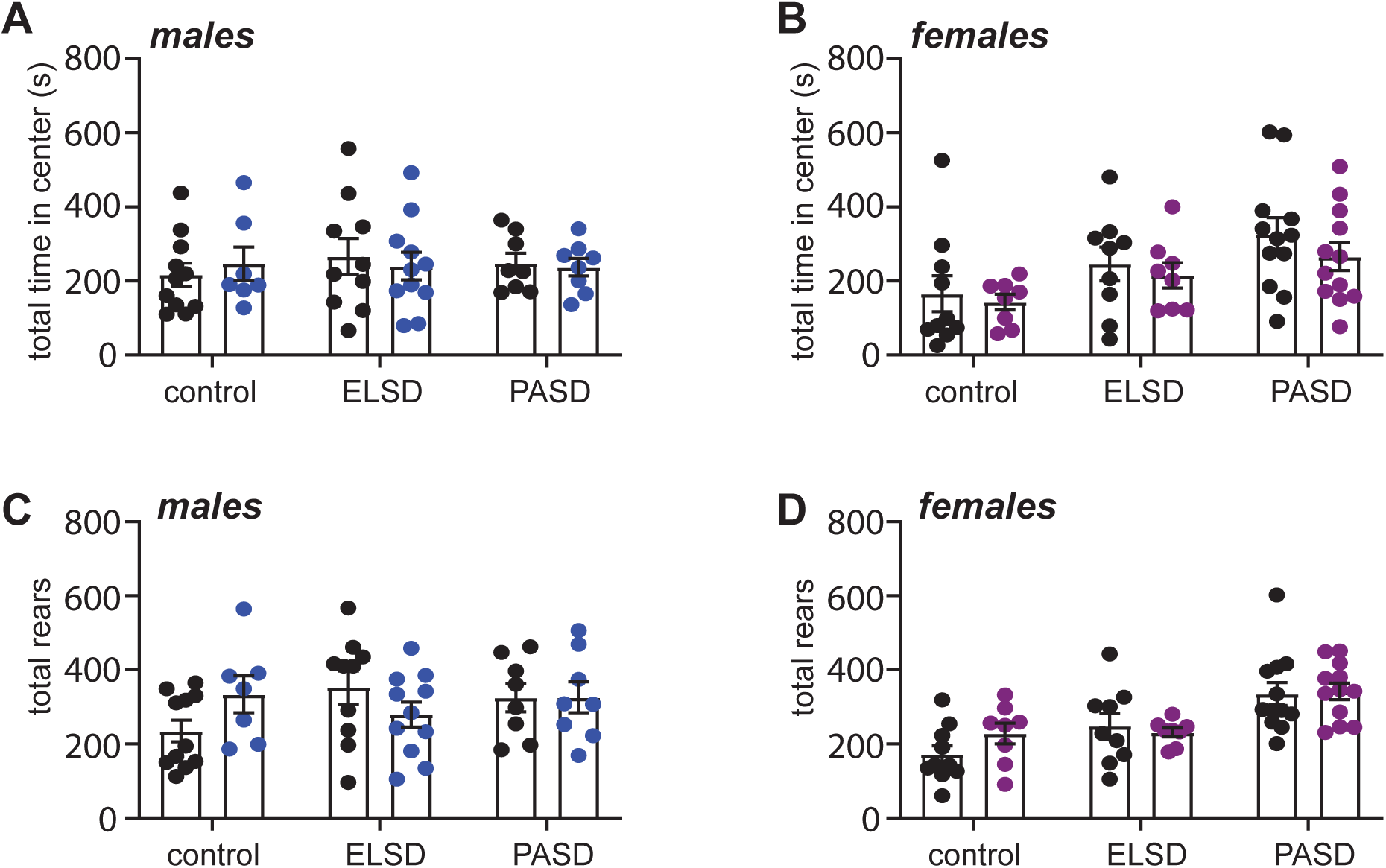
No changes in anxiety-like measures in the open field test. (A-B) no differences observed in total time spent in the center of the open field from control, ELSD, and PASD treated WT and *Shank3*^WT/ΔC^ heterozygous (HET) males (A) and females (B). (C-D) no differences observed in number of rears during the open field test from control, ELSD, and PASD treated males (C) and females (D). (unpaired t-tests with Holm-Šídák correction). N=8-12 per treatment/sex/genotype.

## Notes

### Competing Interest Statement

The authors have declared no competing interest.

### Summary of Updates

We have extended the introduction and discussion sections, in part to incorporate recent advances in the literature. We also include an experimental validation of our early life sleep disruption paradigm.

## References

1. Mazzone L, Postorino V, Siracusano M, Riccioni A, Curatolo P. The Relationship between Sleep Problems, Neurobiological Alterations, Core Symptoms of Autism Spectrum Disorder, and Psychiatric Comorbidities. J Clin Med. 2018;7(5).

2. Missig G, McDougle CJ, Carlezon WA, Jr. Sleep as a translationally-relevant endpoint in studies of autism spectrum disorder (ASD). Neuropsychopharmacology. 2019.

3. Veatch OJ, Sutcliffe JS, Warren ZE, Keenan BT, Potter MH, Malow BA. Shorter sleep duration is associated with social impairment and comorbidities in ASD. Autism Res. 2017;10(7):1221–38.

4. MacDuffie KE, Munson J, Greenson J, Ward TM, Rogers SJ, Dawson G, et al. Sleep Problems and Trajectories of Restricted and Repetitive Behaviors in Children with Neurodevelopmental Disabilities. J Autism Dev Disord. 2020;50(11):3844–56.

5. MacDuffie KE, Shen MD, Dager SR, Styner MA, Kim SH, Paterson S, et al. Sleep Onset Problems and Subcortical Development in Infants Later Diagnosed With Autism Spectrum Disorder. Am J Psychiatry. 2020;177(6):518–25.

6. Verhoeff ME, Blanken LME, Kocevska D, Mileva-Seitz VR, Jaddoe VWV, White T, et al. The bidirectional association between sleep problems and autism spectrum disorder: a population-based cohort study. Mol Autism. 2018;9:8.

7. Jones CE, Opel RA, Kaiser ME, Chau AQ, Quintana JR, Nipper MA, et al. Early-life sleep disruption increases parvalbumin in primary somatosensory cortex and impairs social bonding in prairie voles. Sci Adv. 2019;5(1):eaav5188.

8. Kayser MS, Yue Z, Sehgal A. A critical period of sleep for development of courtship circuitry and behavior in Drosophila. Science. 2014;344(6181):269–74.

9. Jones CE, Chau AQ, Olson RJ, Moore C, Wickham PT, Puranik N, et al. Early life sleep disruption alters glutamate and dendritic spines in prefrontal cortex and impairs cognitive flexibility in prairie voles. Curr Res Neurobiol. 2021;2.

10. Bian WJ, Brewer CL, Kauer JA, de Lecea L. Adolescent sleep shapes social novelty preference in mice. Nat Neurosci. 2022.

11. Roffwarg HP, Muzio JN, Dement WC. Ontogenetic development of the human sleep-dream cycle. Science. 1966;152(3722):604–19.

12. Rensing N, Moy B, Friedman JL, Galindo R, Wong M. Longitudinal analysis of developmental changes in electroencephalography patterns and sleep-wake states of the neonatal mouse. PLoS One. 2018;13(11):e0207031.

13. Nelson AB, Faraguna U, Zoltan JT, Tononi G, Cirelli C. Sleep patterns and homeostatic mechanisms in adolescent mice. Brain Sci. 2013;3(1):318–43.

14. Blumberg MS, Coleman CM, Johnson ED, Shaw C. Developmental divergence of sleep-wake patterns in orexin knockout and wild-type mice. Eur J Neurosci. 2007;25(2):512–8.

15. Penzes P, Cahill ME, Jones KA, VanLeeuwen JE, Woolfrey KM. Dendritic spine pathology in neuropsychiatric disorders. Nat Neurosci. 2011;14(3):285–93.

16. Alcantara S, Ferrer I, Soriano E. Postnatal development of parvalbumin and calbindin D28K immunoreactivities in the cerebral cortex of the rat. Anat Embryol (Berl). 1993;188(1):63–73.

17. del Rio JA, de Lecea L, Ferrer I, Soriano E. The development of parvalbumin-immunoreactivity in the neocortex of the mouse. Brain Res Dev Brain Res. 1994;81(2):247–59.

18. Durand CM, Betancur C, Boeckers TM, Bockmann J, Chaste P, Fauchereau F, et al. Mutations in the gene encoding the synaptic scaffolding protein SHANK3 are associated with autism spectrum disorders. Nat Genet. 2007;39(1):25–7.

19. Costales JL, Kolevzon A. Phelan-McDermid Syndrome and SHANK3: Implications for Treatment. Neurotherapeutics. 2015;12(3):620–30.

20. Bro D, O’Hara R, Primeau M, Hanson-Kahn A, Hallmayer J, Bernstein JA. Sleep Disturbances in Individuals With Phelan-McDermid Syndrome: Correlation With Caregivers’ Sleep Quality and Daytime Functioning. Sleep. 2017;40(2).

21. Smith-Hicks C, Wright D, Kenny A, Stowe RC, McCormack M, Stanfield AC, et al. Sleep Abnormalities in the Synaptopathies-SYNGAP1-Related Intellectual Disability and Phelan-McDermid Syndrome. Brain Sci. 2021;11(9).

22. Speed HE, Kouser M, Xuan Z, Reimers JM, Ochoa CF, Gupta N, et al. Autism-Associated Insertion Mutation (InsG) of Shank3 Exon 21 Causes Impaired Synaptic Transmission and Behavioral Deficits. J Neurosci. 2015;35(26):9648–65.

23. Jaramillo TC, Speed HE, Xuan Z, Reimers JM, Liu S, Powell CM. Altered Striatal Synaptic Function and Abnormal Behaviour in Shank3 Exon4-9 Deletion Mouse Model of Autism. Autism Res. 2016;9(3):350–75.

24. Kouser M, Speed HE, Dewey CM, Reimers JM, Widman AJ, Gupta N, et al. Loss of predominant Shank3 isoforms results in hippocampus-dependent impairments in behavior and synaptic transmission. J Neurosci. 2013;33(47):18448–68.

25. Drapeau E, Riad M, Kajiwara Y, Buxbaum JD. Behavioral Phenotyping of an Improved Mouse Model of Phelan-McDermid Syndrome with a Complete Deletion of the Shank3 Gene. eNeuro. 2018;5(3).

26. Matas E, Maisterrena A, Thabault M, Balado E, Francheteau M, Balbous A, et al. Major motor and gait deficits with sexual dimorphism in a Shank3 mutant mouse model. Mol Autism. 2021;12(1):2.

27. Ingiosi AM, Schoch H, Wintler T, Singletary KG, Righelli D, Roser LG, et al. Shank3 modulates sleep and expression of circadian transcription factors. Elife. 2019;8.

28. Zhou Y, Kaiser T, Monteiro P, Zhang X, Van der Goes MS, Wang D, et al. Mice with Shank3 Mutations Associated with ASD and Schizophrenia Display Both Shared and Distinct Defects. Neuron. 2016;89(1):147–62.

29. Mang GM, Nicod J, Emmenegger Y, Donohue KD, O’Hara BF, Franken P. Evaluation of a piezoelectric system as an alternative to electroencephalogram/ electromyogram recordings in mouse sleep studies. Sleep. 2014;37(8):1383–92.

30. Donohue KD, Medonza DC, Crane ER, O’Hara BF. Assessment of a non-invasive high-throughput classifier for behaviours associated with sleep and wake in mice. Biomed Eng Online. 2008;7:14.

31. Flores AE, Flores JE, Deshpande H, Picazo JA, Xie XS, Franken P, et al. Pattern recognition of sleep in rodents using piezoelectric signals generated by gross body movements. IEEE Trans Biomed Eng. 2007;54(2):225–33.

32. Sinton CM, Kovakkattu D, Friese RS. Validation of a novel method to interrupt sleep in the mouse. J Neurosci Methods. 2009;184(1):71–8.

33. Li Y, Panossian LA, Zhang J, Zhu Y, Zhan G, Chou YT, et al. Effects of chronic sleep fragmentation on wake-active neurons and the hypercapnic arousal response. Sleep. 2014;37(1):51–64.

34. Jimenez JA, Ptacek TS, Tuttle AH, Schmid RS, Moy SS, Simon JM, et al. Chd8 haploinsufficiency impairs early brain development and protein homeostasis later in life. Mol Autism. 2020;11(1):74.

35. Kercmar J, Budefeld T, Grgurevic N, Tobet SA, Majdic G. Adolescent social isolation changes social recognition in adult mice. Behav Brain Res. 2011;216(2):647–51.

36. Kolevzon A, Delaby E, Berry-Kravis E, Buxbaum JD, Betancur C. Neuropsychiatric decompensation in adolescents and adults with Phelan-McDermid syndrome: a systematic review of the literature. Mol Autism. 2019;10:50.

37. Lai MC, Lombardo MV, Auyeung B, Chakrabarti B, Baron-Cohen S. Sex/gender differences and autism: setting the scene for future research. J Am Acad Child Adolesc Psychiatry. 2015;54(1):11–24.

38. Baron-Cohen S, Lombardo MV, Auyeung B, Ashwin E, Chakrabarti B, Knickmeyer R. Why are autism spectrum conditions more prevalent in males? PLoS Biol. 2011;9(6):e1001081.

39. Zhu Y, Fenik P, Zhan G, Xin R, Veasey SC. Degeneration in Arousal Neurons in Chronic Sleep Disruption Modeling Sleep Apnea. Front Neurol. 2015;6:109.

40. Zhu Y, Zhan G, Fenik P, Brandes M, Bell P, Francois N, et al. Chronic Sleep Disruption Advances the Temporal Progression of Tauopathy in P301S Mutant Mice. J Neurosci. 2018;38(48):10255–70.

41. Walsh JJ, Christoffel DJ, Malenka RC. Neural circuits regulating prosocial behaviors. Neuropsychopharmacology. 2022.

42. Moy SS, Nadler JJ, Perez A, Barbaro RP, Johns JM, Magnuson TR, et al. Sociability and preference for social novelty in five inbred strains: an approach to assess autistic-like behavior in mice. Genes Brain Behav. 2004;3(5):287–302.

43. Yang M, Silverman JL, Crawley JN. Automated three-chambered social approach task for mice. Curr Protoc Neurosci. 2011;Chapter 8:Unit 8 26.

44. Lin R, Learman LN, Bangash MA, Melnikova T, Leyder E, Reddy SC, et al. Homer1a regulates Shank3 expression and underlies behavioral vulnerability to stress in a model of Phelan-McDermid syndrome. Cell Rep. 2021;37(7):110014.

45. Ehlen JC, Jones KA, Pinckney L, Gray CL, Burette S, Weinberg RJ, et al. Maternal Ube3a Loss Disrupts Sleep Homeostasis But Leaves Circadian Rhythmicity Largely Intact. J Neurosci. 2015;35(40):13587–98.

46. Johnston MV, Ammanuel S, O’Driscoll C, Wozniak A, Naidu S, Kadam SD. Twenty-four hour quantitative-EEG and in-vivo glutamate biosensor detects activity and circadian rhythm dependent biomarkers of pathogenesis in Mecp2 null mice. Front Syst Neurosci. 2014;8:118.

47. Sare RM, Harkless L, Levine M, Torossian A, Sheeler CA, Smith CB. Deficient Sleep in Mouse Models of Fragile X Syndrome. Front Mol Neurosci. 2017;10:280.

48. Frank MG. Sleep and synaptic plasticity in the developing and adult brain. Curr Top Behav Neurosci. 2015;25:123–49.

49. Zhou Y, Lai CSW, Bai Y, Li W, Zhao R, Yang G, et al. REM sleep promotes experience-dependent dendritic spine elimination in the mouse cortex. Nat Commun. 2020;11(1):4819.

50. Li W, Ma L, Yang G, Gan WB. REM sleep selectively prunes and maintains new synapses in development and learning. Nat Neurosci. 2017;20(3):427–37.

51. Yang G, Gan WB. Sleep contributes to dendritic spine formation and elimination in the developing mouse somatosensory cortex. Dev Neurobiol. 2012;72(11):1391–8.

52. Zoghbi HY. Postnatal neurodevelopmental disorders: meeting at the synapse? Science. 2003;302(5646):826–30.

53. Volk L, Chiu SL, Sharma K, Huganir RL. Glutamate synapses in human cognitive disorders. Annu Rev Neurosci. 2015;38:127–49.

54. Bozdagi O, Sakurai T, Papapetrou D, Wang X, Dickstein DL, Takahashi N, et al. Haploinsufficiency of the autism-associated Shank3 gene leads to deficits in synaptic function, social interaction, and social communication. Mol Autism. 2010;1(1):15.

55. Gai X, Xie HM, Perin JC, Takahashi N, Murphy K, Wenocur AS, et al. Rare structural variation of synapse and neurotransmission genes in autism. Mol Psychiatry. 2012;17(4):402–11.

56. Tatavarty V, Torrado Pacheco A, Groves Kuhnle C, Lin H, Koundinya P, Miska NJ, et al. Autism-Associated Shank3 Is Essential for Homeostatic Compensation in Rodent V1. Neuron. 2020;106(5):769–77 e4.

57. Wood-Downie H, Wong B, Kovshoff H, Mandy W, Hull L, Hadwin JA. Sex/Gender Differences in Camouflaging in Children and Adolescents with Autism. J Autism Dev Disord. 2021;51(4):1353–64.

58. Kreiser NL, White SW. ASD in females: are we overstating the gender difference in diagnosis? Clin Child Fam Psychol Rev. 2014;17(1):67–84.

59. Zhou Y, Sharma J, Ke Q, Landman R, Yuan J, Chen H, et al. Atypical behaviour and connectivity in SHANK3-mutant macaques. Nature. 2019;570(7761):326–31.

60. Wintler T, Schoch H, Frank MG, Peixoto L. Sleep, brain development, and autism spectrum disorders: Insights from animal models. J Neurosci Res. 2020;98(6):1137–49.

